# Genetic variation associated with infection and the environment in the accidental pathogen *Burkholderia pseudomallei*

**DOI:** 10.1101/538793

**Authors:** Claire Chewapreecha, Alison E. Mather, Simon R Harris, Martin Hunt, Matthew T. G. Holden, Chutima Chaichana, Vanaporn Wuthiekanun, Gordon Dougan, Nicholas PJ Day, Direk Limmathurotsakul, Julian Parkhill, Sharon J Peacock

## Abstract

The environmental bacterium *Burkholderia pseudomallei* causes melioidosis, an endemic disease in tropical and sub-tropical countries. This bacterium occupies broad ecological niches ranging from soil, contaminated water, single-cell microbes and plants to human infection. Here, we performed genome-wide association studies for genetic determinants of environmental and human adaptation using a combined dataset of 1,011 whole genome sequences of *B. pseudomallei* from Northeast Thailand (discovery cohort, number of clinical isolates = 325, environmental isolates =428) and Australia (validation cohort, number of clinical isolates = 185, environmental isolates = 73), representing two major global disease hotspots. With these data, we (i) identified 47 genes from 26 distinct loci associated with clinical or environmental isolates from Thailand and replicated 13 loci in an independent Australian cohort; (ii) outlined the selective pressures on the genetic loci (dN/dS) and the frequency at which they had been gained or lost throughout their evolutionary history, reflecting the bacterial adaptability to a wide range of ecological niches; and (iii) highlighted loci implicated in human disease.

## Results

*Burkholderia pseudomallei* is an environmental Gram-negative bacterium and the cause of melioidosis, a serious infectious disease that affects an estimated 165,000 people and causes around 89,000 deaths per year globally^1^. The bacterium has a broad range of ecological niches, and may be isolated from soil, surface water, amoebae, plants, animals, and humans in many tropical and sub-tropical regions.^2-4^ Human infection results from environmental exposure associated with inoculation, ingestion or inhalation of the bacterium, with increasing risk of acquisition for people with predisposing health conditions or activities that increase exposure to soil or water such as rice farming or drinking untreated water^5^. Infection can be acute, chronic, latent or cleared^6^, with rare cases of human-to-human transmission being reported^7,8^. Antibody responses to *B. pseudomallei* can be found in healthy individuals living in endemic areas in the absence of clinical symptoms^9,10^, suggesting that the majority of the exposure is harmless or results in sub-clinical infection. *B. pseudomallei* can be found in the stools of infected humans^11^ and experimental murine models.^12^ This provides a potential mechanism for human-to-environmental transmission and the possibility of repeated passage through the human host. Serial passage of *Burkholderia cenocepacia* in a long-term chronic airway infection model in mice has been shown to increase bacterial fitness ^13^. Based on this observation, the natural passage of *Burkholderia pseudomallei* through humans, animals or its natural predators such as soil amoebae might have enhanced and maintained selection pressure for pathogenicity in a subset of the population. This potentially results in heterogeneity of bacterial virulence, as evidenced by marked variations in severity and pathogenicity in mice challenged by different *B. pseudomallei* strains^14-16^. *B. pseudomallei* has a large and highly variable accessory genome across the species^17-19^. While the core genome may be sufficient for strain survival, it is possible that specific bacterial genes, gene variants or their combinations may confer additional advantages for survival and replication in specific niches including human infection, or particular environmental conditions.

Here, we sought evidence for bacterial genetic factors associated with human disease and environmental adaptation using two independent datasets from major melioidosis hotspots in Thailand (325 clinical and 428 environmental isolates), and Australia^18-23^ (102 clinical and 63 environmental isolates). These were used as a discovery and validation dataset, respectively (Supplementary Fig. 1). The discovery dataset was originally cultured from patients presenting to Sunpasitthiprasong hospital in Ubon Ratchathani, Northeast Thailand between 2010-2011, together with their residential water sources^5^ (see methods for details). Cases were patients with melioidosis and controls were patients without melioidosis admitted to the hospital during the same period. We sequenced 325 clinical isolates from 324 patients with melioidosis.

### Not all environmental exposure leads to melioidosis infection

The household sampling structure in the discovery cohort allowed us to examine the minimum *B. pseudomallei* diversity to which each patient was potentially exposed (up to 10 isolates per patient household) and compare this with their infecting isolate (1 isolate per patient). Consumption of untreated household water was common (21/27 cases and 17/21 of controls), and associated with a higher risk of melioidosis^5^.

We measured the genetic diversity of water isolates and their matched clinical isolates against other isolates from a circumscribed area of Northeast Thailand using three methods: multi-locus sequence typing (MLST)^24,25^, pan-genome analysis^26^, and using a kmer approach^27^. We excluded households with < 2 environmental isolates, leaving samples from 25 cases and 19 controls. Each water sample contained a mean of 2.5 *B. pseudomallei* sequence types (STs) (range 1 – 6 STs), with no difference in bacterial diversity per household between cases and controls by MLST (Mann-Whitney U test, p-value 0.785), pair-wise distance of core gene single nucleotide polymorphisms (SNPs) (Mann-Whitney U test, p-value =0.230, Supplementary Fig. 2), or pair-wise kmer distance (Mann-Whitney U test, p-value = 0.778). A subgroup analysis based on diabetic status did not observe any differences between diabetics and non-diabetics.

We next quantified how much of the exposure had developed into clinical melioidosis. MLST analysis identified shared STs between water and clinical isolates in 3/27 cases (A-175, A-266 and A-330) with no difference in ST distribution between environmental and clinical sources (average Simpson Diversity index: D_clin_= 0.959, D_env_ = 0.953; Pielou’s Evenness: J_clin_ = 0.956, J_env_ = 0.968; ANOSIM Benjamini-Hochberg adjusted p-values > 0.100 for 100 permutations), which is consistent with previous studies^28,29^. However, none of the isolates from water samples and their matched clinical isolates were closely related at the whole-genome level (Figure 1 a, Supplementary Fig. 2). The closest genetic match between a clinical and water isolate pair was found in patient A-175 who reported consuming untreated water from their domestic sources prior to presenting with melioidosis. This pair were 483 SNPs (1.07 x 10^-4^ SNPs per site) different on core genome comparison, and had 57 and 38 genes uniquely present in the clinical and environmental isolate, respectively. Assuming a constant combined substitution^30^ and recombination^31^ rate detected in *B. pseudomallei* (~ 8.2 x 10^-7^ SNPs per site per year), this is consistent with ~130 years of divergence between the clinical and its closest household water isolate for A-175. This effectively rules out the chance of the environmental isolates being immediately parental to the infecting isolate. It is possible that the infecting isolate was acquired elsewhere, that it represented a minority population in water that was not detected in this study, or that the diversity varies over time. Nevertheless, these results still argue for a selective process or bottleneck during acquisition, where not all isolates the patients are exposed to lead to infection (albeit the bottleneck stringency may be influenced by routes of acquisition and amount of exposure, as seen in ^32^).

**Fig. 1.**
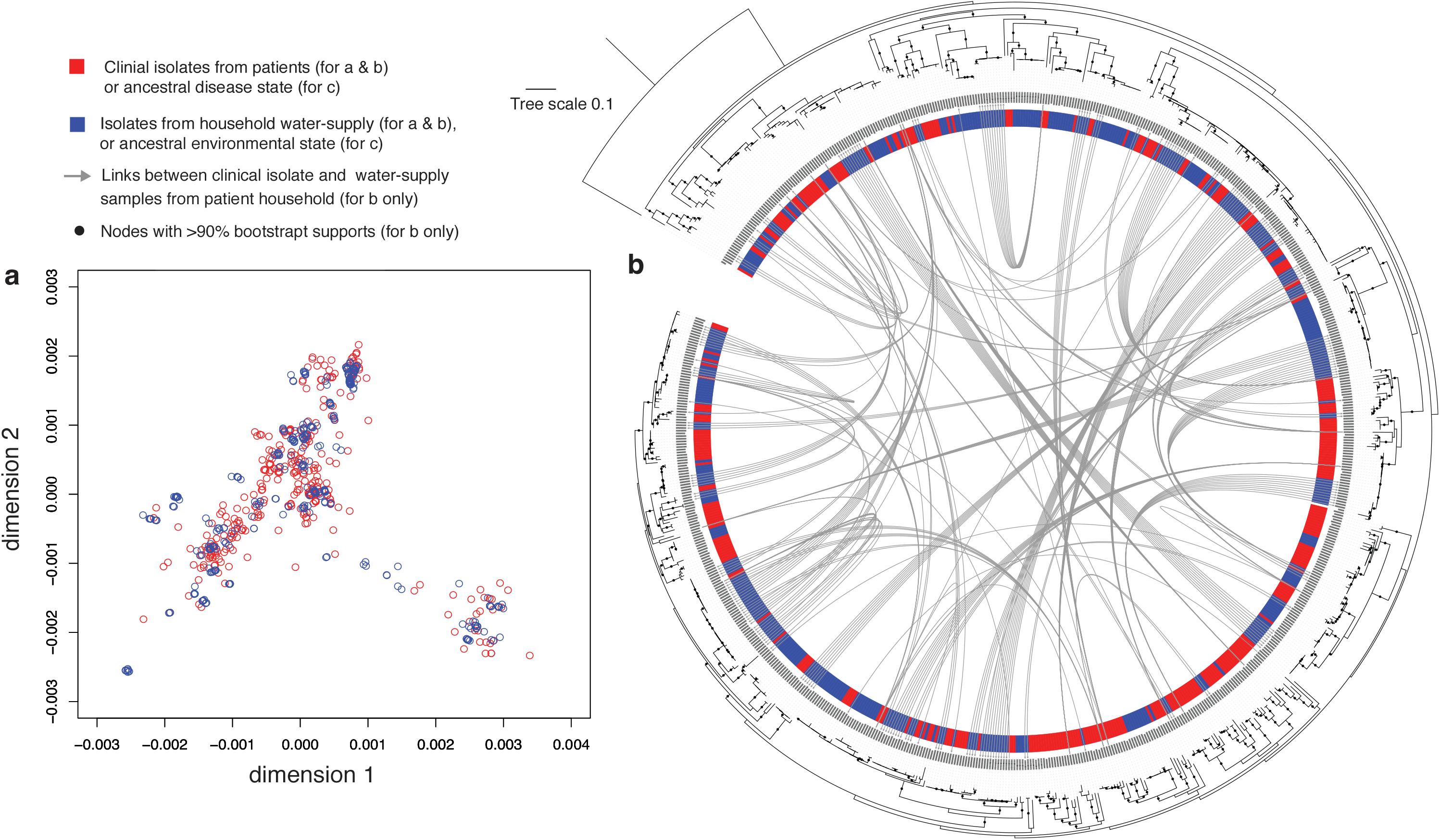
Population structure and phylogeny of B. pseudomallei isolated from patients and their household water supplies in northeast Thailand. a) Multi-dimensional scaling based on the two dimensions that best explained data variability. b) Maximum likelihood phylogeny generated from core gene SNPs in northeast Thailand isolates rooted on an Australasian outgroup Bp668. Nodes with bootstrap support of >90 are shown by black dots, and the coloured ring shows the isolate source. The grey arches represent an analysis of 27 cases who had a clinical isolates and up to 10 isolates cultured from their water supply, showing the connections for isolates from each case.

### Population structure of clinical and environmental isolates

We next evaluated the genetic content of the clinical and environmental isolates in the discovery cohort by including a further 298 clinical isolates from patients admitted to the same hospital (in total, 325 clinical and 428 environmental isolates).

Unlike many pathogens where isolates associated with disease contain substantially fewer genes^33,34^, we observed a similar number of genes per genome (two-sided Mann-Whitney U test, p-value = 0.312) in clinical and environmental isolate collections. The population structure of clinical and environmental isolates was defined using a phylogenetic tree generated from core genes and multidimensional scaling (MDS) generated from kmers (Fig. 1b & c). These results indicated that clinical and environmental isolates were inter-mixed and co-circulating. We next tested for potential genetic signals that were associated with infection or the environment by estimating the correlation between bacterial phylogeny and distribution of source of isolation on the tree using Pagel’s λ^35^ and Bloomberg’s K^36^. The distribution of “disease” and “environmental” origins was not random after correcting for sampling bias (Supplementary Fig. 3), indicating that there may be separable environmental and clinical clades either at deep or shallow nodes^37^. This could reflect the presence of bacterial determinants that mediate survival in human or environmental niches.

### Genetic determinants associated with disease and environmental isolates

We applied two complementary genome-wide association (GWAS) methods (a kmer-based^27^ and a pan-genome based^38^) to the 325 clinical and 428 environmental isolates, which were controlled for population stratification (see methods). We note that our “environmental” isolates could be capable of causing disease. However, this should only reduce our power to detect association, not introduce false positive associations. Of 24,856,071 kmers used to define the population, 38,813 (0.156 %) were associated with “disease” or “environmental” origin. These were mapped onto the pan-genome to identify potential genes, resulting in 365 “disease-associated” or “environmental-associated” genes (Fig. 2). The pan-genome based GWAS analysis identified 675 disease-associated or environment-associated genes. Comparison of output from the two methods showed that 47 genes were detected by both (38 disease-associated and 9 environmental-associated genes), which account for 0.3% of the pan-genome. Based on the size of operons of transcription fragments reported in Ooi *et al*.^39^, we grouped these genes into 26 loci (Supplementary Fig. 4). These 47 genes were evaluated in an independent dataset from Australia (clinical isolates = 185, environmental isolates = 73), which showed that 13 gene clusters (27.7%) could be replicated (two-sided Fisher’s exact test, FDR < 0.01). The fact that isolates from Australia and Southeast Asia represent distinct phylogenetic clades^19,40,41^ is consistent with parallel evolution for a proportion of the disease-associated and environment-associated genes.

**Fig. 2.**
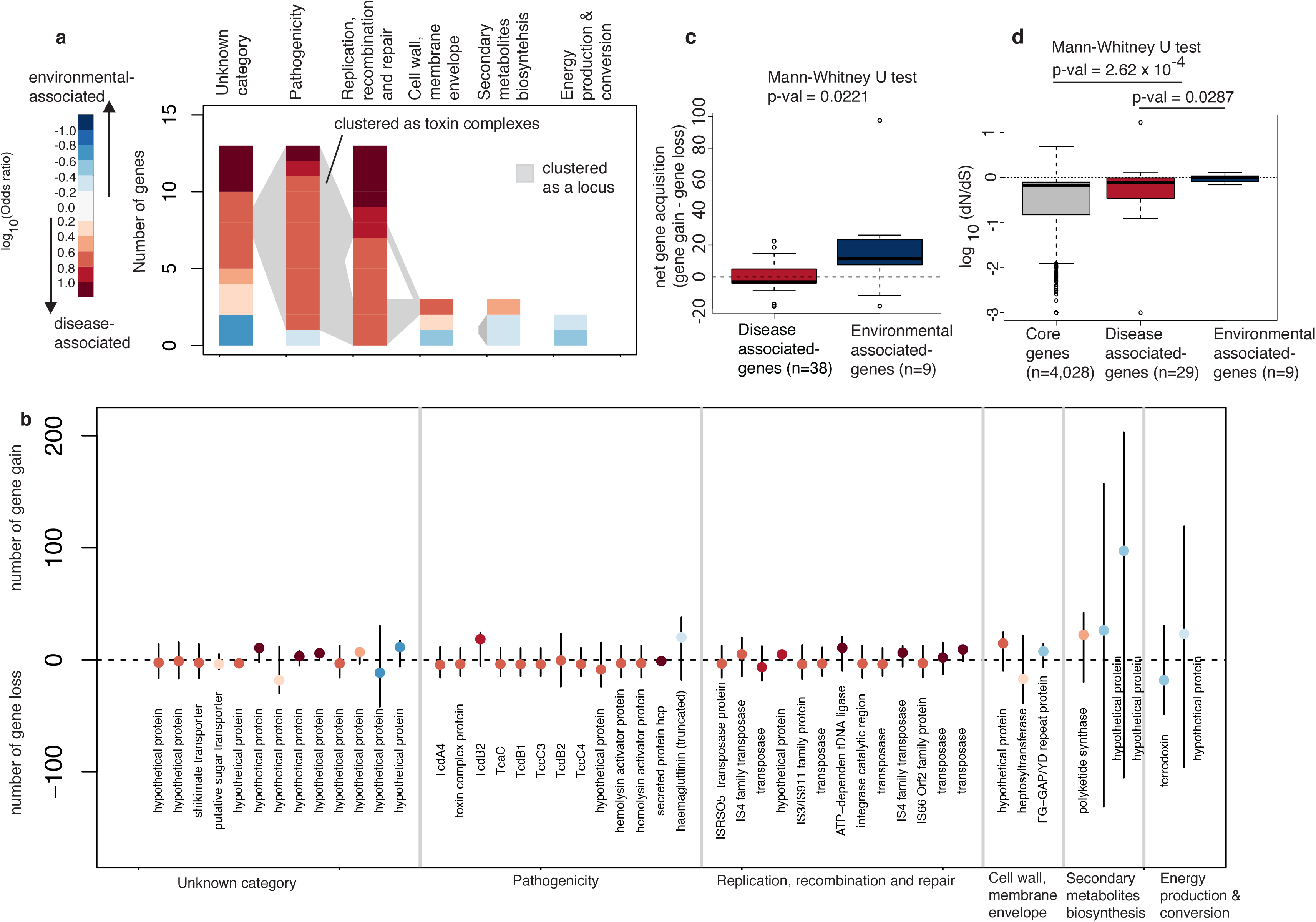
*B. pseudomallei* disease- and environmental-associated genes. (a) Bar charts summarise the frequency of disease-or environment-associated genes by functional category. The plots are ranked by categorical gene frequency from unknown category (13), potential roles in pathogenicity (13), replication, recombination and repair (13), cell wall membrane envelope biogenesis (3), secondary metabolite biosynthesis (3), and energy production and conservation (2). Colour shading for each gene reflects the directionality of gene association in the discovery dataset on the log10(Odds ratio) scale with red and blue denoting disease and environmental association, respectively. (b) Frequency of gene gain and loss throughout the phylogeny with the net gain or loss summarised as dot in the middle of each bar. The colour of the dot indicates the disease- and environmental association using the same colour scheme as (a). (c) Boxplot comparing the net gene gain and loss events for disease-associated genes (red) and environmental associated genes (blue). (d) The dN/dS of core genes (grey) disease-associated genes (red) and environmental-associated genes (blue) are plotted on a log 10 scale. For (c) and (d) boxplots summarise the distribution of data based on first quantile, median, and third quantile. Two-sided Mann-Whitney U test was used to compare categorical observation in (c) and (d).

Functional enrichment analyses of the 47 gene clusters in the discovery cohort showed an elevated frequency of the term “Pathogenesis” and “Replication, recombination and repair” (One-sided Fisher’s exact test p-value 2.30 x 10^-7^, and 2.08 x 10^-12^ respectively). The former may allow the bacterium to compete in specific environmental niches or survive inside single-cell or multicellular organisms during infection. Genes annotated with the term “Replication, recombination and repair” largely comprise transposons that may act as markers or remnant elements for horizontally transfered genes, or may inactivate gene function. Notably, 8 of 26 loci consist of IS, transposons and integrase, which highlights the significance of transposable elements in modulating bacterial genomes.

### Selection pressures maintaining disease- and environmental-associated genes

We explored whether the 38 disease-associated and 9 environmental-associated genes were under selective pressure by calculating the ratio of non-synonymous to synonymous substitutions (dN/dS). The average for both groups were below 1, but the ratio was significantly higher for environmental-associated compared with disease-associated genes (Fig. 2b, Mann-Whitney U test = 0.0287), with both environment- and disease-associated genes having higher ratio than core genes (Mann-Whitney U test = 2.62 x 10^-4^). Despite a small number of genes being compared, this suggests that the subset of genes in the environment-associated genes may be under reduced purifying selection, or elevated diversifying selection, compared to disease-associated genes and core genes, with disease-associated genes also having a stronger signal than core genes. We further quantified the number of times each cluster was acquired or lost from phylogenetic branches (Fig 2b & c). Stochastic mapping of the presence of each disease-or environment-associated cluster estimated that 45/47 genes were both acquired and lost multiple times. One possible reason for multiple gain-and-loss events is a constant change in niches, which could include switching between extra- and intracellular lifestyles. However, the result should be interpreted with care as internal nodes on the phylogenetic tree of many recombinogenic species including *B. pseudomallei* are not well resolved.

### Examples of disease-and environmental associated genes

Many of the disease-associated loci contained genes that encoded biologically plausible or known virulence determinants. One example was a large toxin complex *(tcdB, tcdA, tccC* and hemolysin activator *fhaC)* encoded by a locus of up to 69.7 kb, which was identified in the discovery dataset (Supplementary Fig. 5). This locus has not been characterised in *B. pseudomallei* but homologues exist in diverse bacterial species including *Pseudomonas*, *Yersinia* and *Photorhabdus^42,43^*. The latter is an insect pathogen, experimental characterisation of which has demonstrated that *tccC* has enzymatic activity^44^ and that *tcdA* and *tcdB* facilitate the translocation of the toxin into host cells^43^. These toxin genes were flanked in *B. pseudomallei* by several integrases and transposases from families including IS2, IS3/IS911, IS4, IS66, IS166, IS407, IS111A/IS1328/IS1533, and IS1478, indicative of a mobile genetic element origin. Except for *tcdB2*, genes in this locus had been acquired up to 15 and lost 24 times in the Thai dataset. A greater net loss may suggest a fitness cost associated with this locus.

An example of environmental-associated loci is a truncated variant of filamentous hemagglutinin *(fha*), a known adhesin and immunomodulator across different bacterial species. In *B. pseudomallei,* the number of *fha* genes varies between isolates, and different combinations of *fha* genes have been observed with patients infected by *B. pseudomallei*, with a specific *fha* variant reported to have increased risk of infection associated with positive blood cultures^45^. While our kmer approach identified disease-associated signals from haemaglutination activity domains on this gene, our pan-genome approach detected environmental-associated signals from a truncated form of this gene (Supplementary Fig. 6). A closer inspection highlighted a truncation caused by a premature stop codon upstream of the haemaglutinin repeat domains, which might disrupt gene function.

## Discussion

Our analyses have identified *B. pseudomallei* gene clusters that distinguish between isolates associated with human disease *vs* environmental isolates. These genes have arisen repeatedly in different populations with distinct phylogeography, demonstrating robustness of the findings from the Thai discovery dataset. Many of these genes are under relaxed purifying selection and have been gained or lost multiple times throughout evolutionary history, implying that there may be several niches to which this opportunistic bacterium is adapted. This includes environmental and other eukaryotic hosts^3,46,47^, the latter potentially providing genetic pre-adaptation for invasion and survival in the human host. Our catalogue of disease-associated genes adds new information for future studies of disease pathogenesis including an evolutionary dimension, providing opportunities to further explore host-pathogen interactions.

## Methods

### Bacterial isolates

Two bacterial collections were used to create independent discovery and validation datasets. These originated from distinct regions where melioidosis is highly endemic – northeast Thailand (753 isolates), and northern Australia^18-22^ (165 isolates). The discovery dataset was drawn from a study of the activities of daily living associated with melioidosis, which was conducted at Sunpasitthiprasong (formerly Sappasithiprasong) hospital in Ubon Ratchathani, Northeast Thailand between 2010-2011^5^. In brief, 330 cases of culture-proven melioidosis and 513 control patients with non-infectious conditions were recruited^5^. Residential drinking water was collected and cultured for *B. pseudomallei* from cases and controls who lived within 100 km of the hospital. *B. pseudomallei* was isolated from 12% of borehole and tap water samples, and 4% of well water samples. Multiple colonies were picked and individually saved from each water sample. Consumption of untreated water was common (85% of cases and 72% of controls) and associated with a higher risk of melioidosis.^5^ For the purposes of the study described here, we sequenced 325 *B. pseudomallei* isolates from 324 cases, and 428 *B. pseudomallei* colonies (isolates) from 48 water samples (including samples from 27 melioidosis patients) (Supplementary Fig. 1). We assumed that isolates from water were a fair representation of environmental isolates. The validation dataset consisted of whole genome sequence data for 258 *B. pseudomallei* isolated in Australia, which were downloaded from the NCBI database. These isolates have been described previously^18-23^. In brief, isolates were from patients with melioidosis (n=185) and the environment (n=73). The temporal and spatial distribution of isolates in this dataset is summarised in Supplementary Fig. 1.

### Whole-genome sequencing

DNA was extracted from the 753 Thai *B. pseudomallei* isolates as described in ^48^. DNA libraries were prepared according to the Illumina protocol and sequenced on an Illumina HiSeq2000 with 100-cycle paired-end runs to give a mean coverage of 84 reads per nucleotide. Sequencing of clinical and environmental isolates was done at the same time on the same platform. Taxonomic identity was assigned using Kraken^49^.

### Genome assembly and pan-genome analysis

New assemblies were performed as described in ^50^ to give a median of 97 contigs (min = 61 contigs, max = 259 contigs), and median length of 7,114,540 bp (min=6,884,381 bp, max= 7,404,549 bp). All study genomes were annotated using Prokka^51^. A predicted median of 5,936 coding sequences were assigned onto each genome (min = 5,762, max = 6,264), which falls within the range of published reference genomes^46,52^. Roary^26^ was used to calculate the pan-genome for the discovery dataset together with the two reference *B. pseudomallei* genomes (K96243 from Thailand and Bp668 from Australia). The inclusion of the well-characterized Thai reference K96243 served as the quality control for the pan-genome analysis, and the Australian reference Bp668 served as an outgroup to root the phylogeny in a subsequent analysis. An all-against-all BLASTP comparison at 92% sequence identity was used as described in^19^. Genes were defined as core if present in ≥99% of isolates. This led to 4,322 and 10,718 genes being classified as core and accessory, respectively. The number of core genes identified fell within the range described previously^18^.

### Estimating the population structure using multi-dimensional scaling

The population structure of the 753 Thai isolates was estimated from sequence assemblies using Mash v. 1.1.1^53^, which captures information from intergenic regions, core and accessory genes. Assemblies were shredded into their constituent kmers. The pairwise distance between assemblies was estimated and computed into the 753 x 753 matrix. Metric multi-dimensional scaling (MDS) was performed using R cmdscale to project the population structure into n-1 coordinates. The top three coordinates were used to control for GWAS population stratification.

### Estimating genome and gene evolution with phylogenetic trees

The population structure was also estimated using a phylogenetic tree based on SNPs in the core genome. Single-copy core genes from 753 isolates, K96243 and Bp668 were concatenated and aligned using Mafft v7.205^54^, followed by manual inspection using SeaView^55^. This comprised 4,322 genes, representing on average 73% of genes in individual genomes. Single nucleotide substitutions in the alignment were called using the methods described by Page *et al.* 2016^56^, resulting in SNPs. A maximum-likelihood phylogeny was constructed with RAxML HPC v.8.2.8^57^ using a general time reversible nucleotide substitution model with four gamma categories for rate heterogeneity and 100 bootstrap support. The overall phylogeny had 58.4 % of external and internal nodes showing >70% bootstrap support. The phylogeny has been archived at: ftp/pub/pub/project/pathogens/Burkholderia/RAxML_bipartitions.Ubon.tree. Maximum-likelihood phylogenies were used to examine specific disease-associated clusters by concatenating and aligning genes using Mafft v7.205^54^, with truncated genes manually checked. A maximum-likelihood phylogeny was constructed as above with 100 bootstrap support and rooted on an Australian gene homologue.

### Estimation of phylogenetic signals

Pagel’s λ^35,58^ was used to assess phylogenetic signal in the Thai dataset. Origin of isolation (clinical or environmental) was reconstructed onto the tree using fitDiscrete from the R package Geiger^59^. We compared the model fit of the tree using log-likelihood of the untransformed maximum likelihood tree against the model where the tree was transformed to a polytomy or partially transformed trees (internal branches were multiplied by λ = 0, 0.1, 0.2, 0.3, 0.4, 0.5, 0.6, 0.7, 0.8, and 0.9, Supplementary Fig. 3). We also reconstructed randomised origin of isolation (clinical or environmental, 100 permutations) onto the tree and compared log-likelihood scores obtained from reconstruction with the actual origin versus randomised origins. Each reconstruction was performed with 3 different simulation models (equal rate transition (ER), a symmetric mode (SYM), and all rates different matrix (ARD)). We also tested phylogenetic signals using Bloomberg K^36^ using R package picante v. 1.7 ^60^ to provide further evidence of potential signals. To investigate whether results were affected by sampling bias caused by using multiple isolates from the same environmental sample, we sub-sampled the dataset to have a single environmental isolate from each of 48 households and an equal number of clinical isolates and resampled this 100 times. We estimated the log-likelihood of the fit of disease *vs* environmental origins on the tree with real and randomised data.

### Detecting kmers associated with clinical and environmental isolates

Two separate GWAS were performed to screen kmers for associations with source of isolation (clinical or environmental) using the 753 Thai genomes. Assemblies were shredded into kmers of 9-100 bases, resulting in 24,856,071 kmers. All kmers occurring in more than one assembly were counted using fsm-lite (https://github.com/nvalimak/fsm-lite) as described in ^27^, and filtered to retain kmers that appeared in 5%-95% of samples. This is equivalent to the recommended 5% minor allele frequency cut-off and reduced the data to 24,555,746 kmers. Further stringency was added by testing kmer association using the χ^2^-test (1 d.f.), in which only those with a p-value <10^-5^ were retained. This has been shown previously through simulations to select for true positive associations^27^, and reduces the number of kmers linked to a lineage but unrelated to true phenotypes. This step reduced the kmers to 300,325 kmers. Seer^27^ was then used to fit a logistic curve to binary data (clinical or environmental) for each kmer. The first three principal components calculated from metric dimensional scaling were used as covariates to control for bacterial population structure. This resulted in 37,104 kmers associated with clinical isolates, and 1,709 kmers associated with environmental isolates. All kmers were mapped to the K96243 reference^52^ and the raw assemblies of 753 isolates using BLAT v. 35^61^ to identify the relevant genes and gene variants. To allowed for mapping of low complexity kmers (length 10-26 base pairs), the following parameters were used: blat –minMatch=1 –tileSize=8 –minScore=10. Only identical hits were retained. Any predicted coding sequences (CDS) with more than 2 kmer hits were pooled and collectively termed “disease-associated” genes or “environment-associated” genes.

### Determination of genes that distinguish clinical and environmental isolates

As a complementary approach, we performed a pan-genome based GWAS using Scoary^38^ on the Thai dataset. Two separate GWAS were performed to find genes associated with source of isolation (clinical or environmental) while correcting for population structure using the phylogenetic tree. Benjamini-Hochberg’s method was used to correct for multiple comparisons. Disease-associated or environment-associated genes with a Benjamini-Hochberg adjusted p-value < 0.01 were reported and compared for consistency with genes identified by the kmer-based GWAS.

### Validation of genes associated with clinical and environmental source

Disease-associated or environment-associated genes that were identified by both the kmer-based and pan-genome based methods (n = 47) were validated in an independent dataset from Australia. Where genome assemblies were available, genes were validated by searching for Australian orthologues using BLAT v. 35^61^ allowing for 92% identity, the same cut-off used in the pan-genome analysis. Where short reads were available^23^, ARIBA^62^ was employed to perform local assembly and mapping to check for the presence or absence of genes. The sequence identity threshold (-- cdhit_min_id 92) was adjusted to 92% for consistency. Gene distribution across Australian clinical and environmental isolates was tested using two-sided Fisher’s exact test with a Benjamini-Hochberg adjusted p-value.

### Functional category annotation and enrichment in pan-genome and disease-associated and environment-associated genes

Gene ontology (GO) describing biological process, molecular function and cellular compartment was assigned to each gene in the pan-genome using InterProscan v5.21-60.0^63^. Not all genes matched the GO database. As of October 2018, 38.2% genes had GO terms assigned. A given gene could be associated with multiple GO terms (mean~2.85, min=1, max =14), and when this occurred a parent GO term was used to represent child GO terms. Comparison of GO terms in disease versus environmental isolates, and their enrichment among disease-associated clusters versus expectation based on the reference genome K96243 was performed using one-sided Fisher’s exact test with all GO terms, with a Benjamini-Hochberg adjusted p-value.

Disease-associated clusters were also annotated with Orthologous Groups of Proteins (COG^64^) and pathway maps (KEGG^65^ and MetaCyc^66^) to determine putative function. As of October 2018, COG, KEGG and MetaCyc could be assigned to 78.04%, 9.79% and 7.72% of disease-associated genes, respectively. Information on protein domains was sourced from the Conserved Domain Database (CDD)^67^.

### Measuring selection pressure acting on disease- and environment-associated genes

The ratio of non-synonymous to synonymous substitutions (dN/dS or Ka/Ks) was calculated using the KaKs calculator^68^. Alignments of core, disease-associated and environment-associated genes were extracted from the pan-genome^26^. The test rejected neutrality (H_0_ dN/dS = 1, Fisher’s exact test p-value < 0.05) in 4,028 core, 28 disease-associated and 9 environment-associated genes. A non-parametric Mann Whitney U test was used to investigate any departures in the mean of dN/dS for genes associated with disease, the environment, and core.

### Estimating gain and loss of disease-associated and environment-associated genes

Gain or loss of disease-associated and environment-associated genes through evolutionary history was quantified using make.simmap from the R package Phytools v 0.6-44^69^ with 1,000 simulations. We first compared likelihood scores for the presence or absence of each gene across the phylogeny with the three different models (AR, ER and SYM). ER was the best fit model in our dataset and was selected. Only genes with a frequency > 0.01 were included in the analyses.

### Statistics and visualisation

Visualisation of phylogenetic trees and statistical analyses was performed in R, Phandango^70^, and FigTree v 1.4.2 (http://tree.bio.ed.ac.uk/software/figtree/).

---

**Supplementary Fig. 1.**
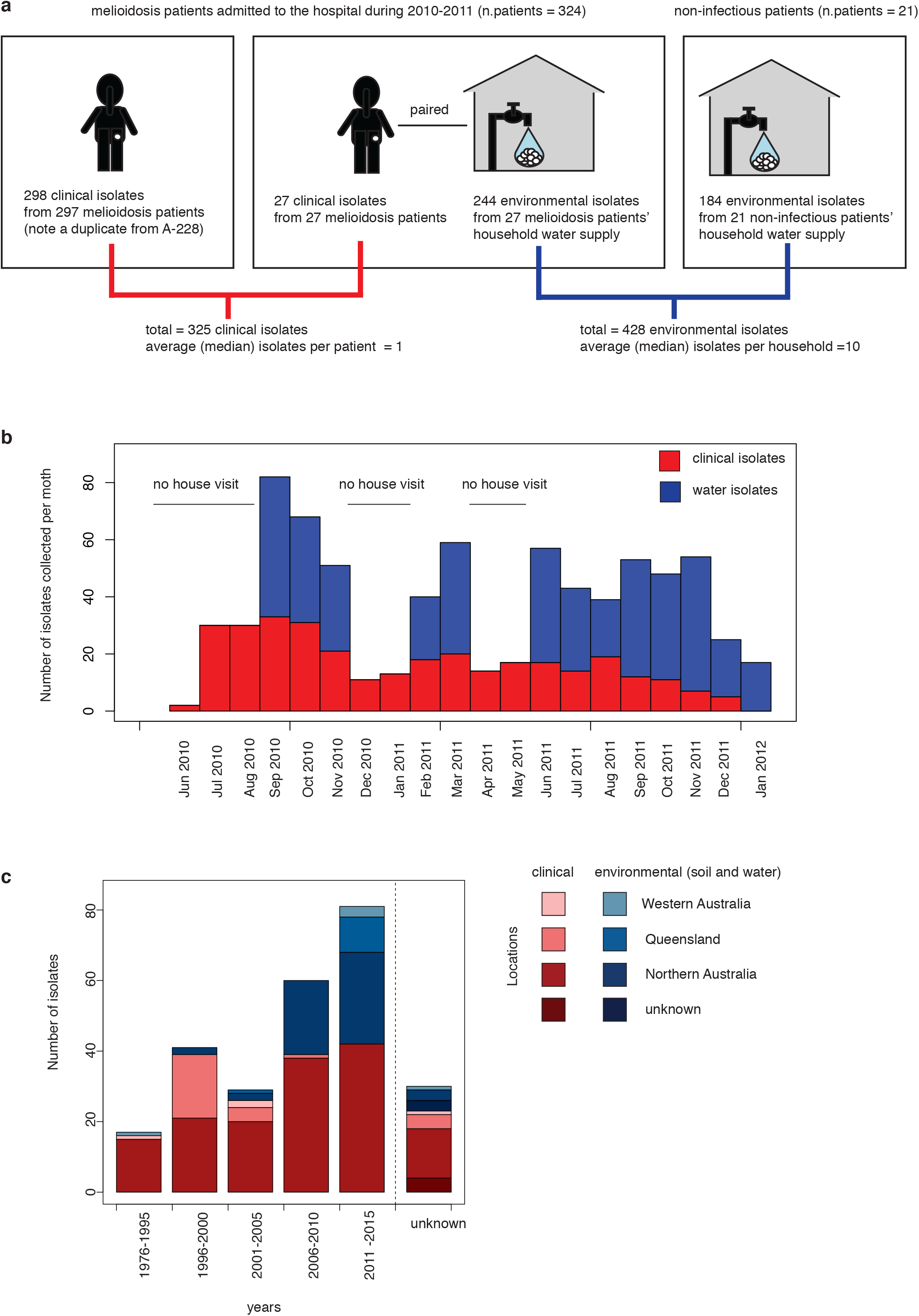
Sampling framework for *B. pseudomallei* isolates from the case control study. (a) The chart shows the number of clinical and environmental isolates from patients and/or household water supplies of cases (patients with melioidosis) and controls (patients with non-infectious conditions admitted during the same period). (b) Temporal distribution of environmental and disease isolates in the discovery dataset collected from June 2010 – January 2012. With the exception of months with no house visits, the number of monthly clinical and environmental samples collected were positively correlated (linear regression, adjusted R-square = 0.259, p-value = 0.026). (c) Spatial and temporal distribution of environmental and disease isolates in the validation dataset from the public database.

**Supplementary Fig. 2.**
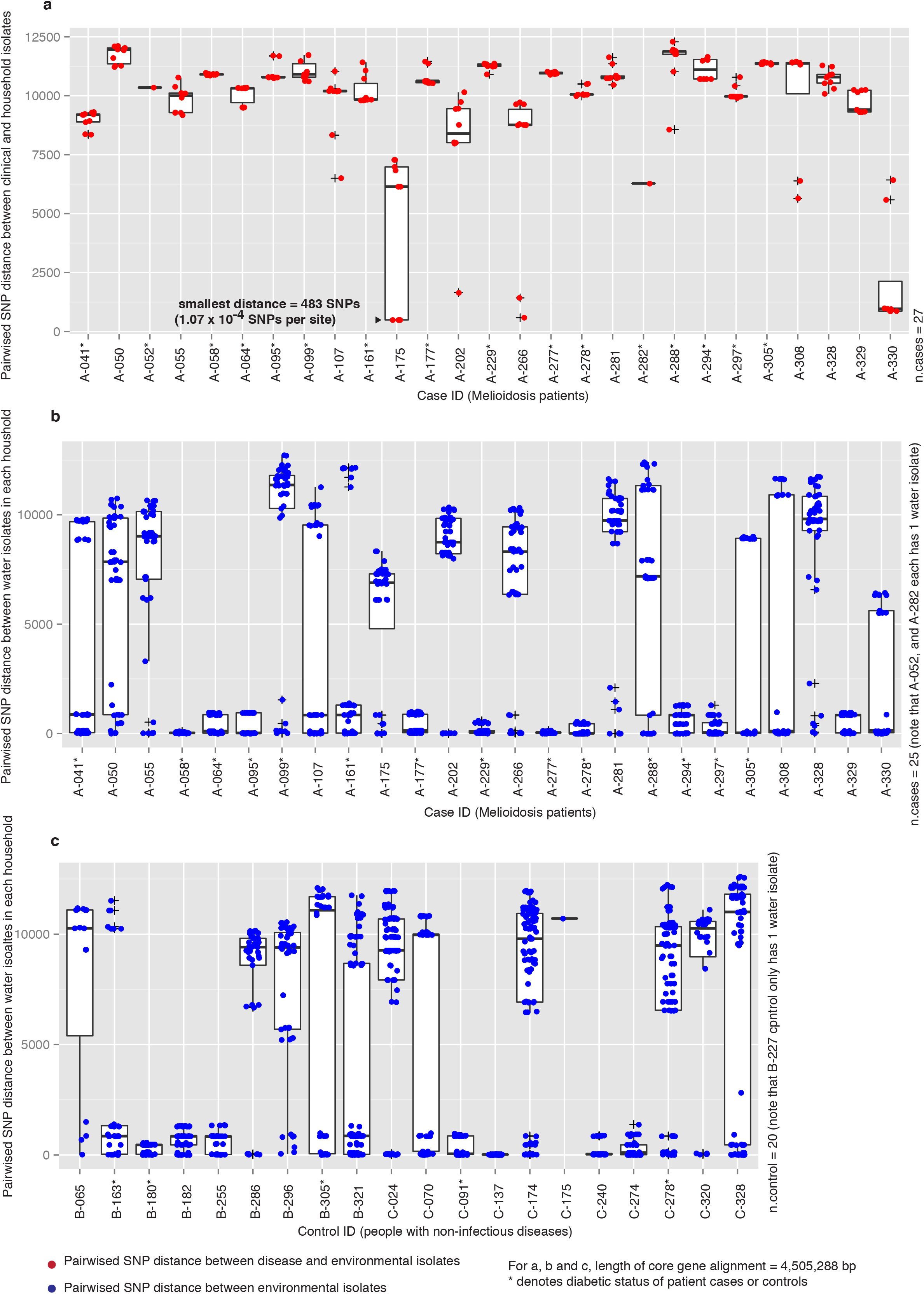
Epidemiological analyses of clinical and/or environmental isolates collected from each patient and his or her household for 27 patients. a) For a case with melioidosis where both clinical and environmental isolates were available, the pairwise SNP distance was computed. The red boxplot shows the pairwise SNP distance between clinical and environmental isolates collected from the same patient-household water supply. b) The blue histogram shows the pairwise SNP distance within environmental isolates. Up to 10 environmental isolates were available for each clinical isolate. c) For control patients (with non-infectious disease) where only environmental isolates were available. Patients with diabetes mellitus are marked with asterisks.

**Supplementary Fig. 3.**
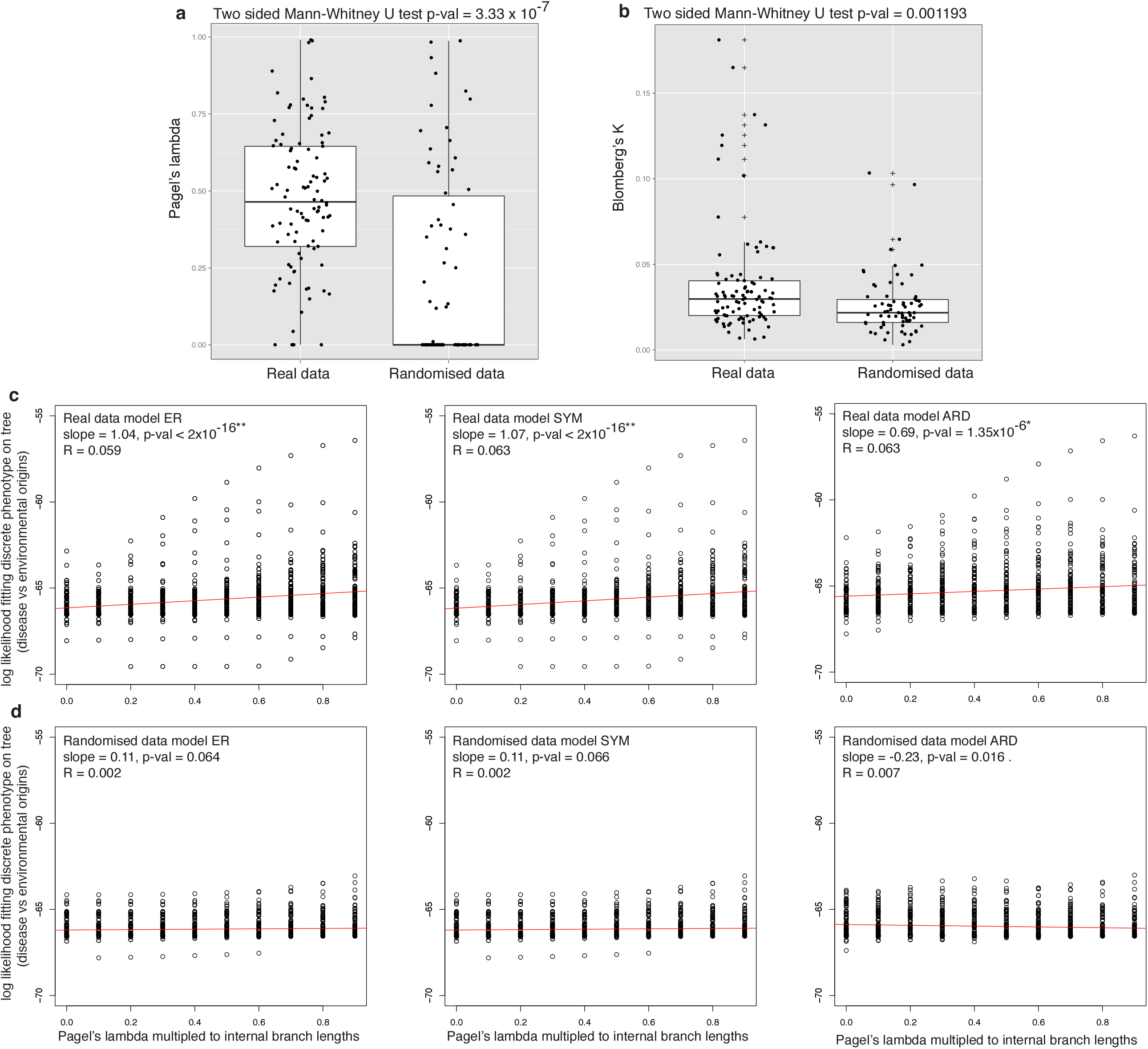
Measurement of phylogenetic signals correlated with bacterial clinical and environmental origins using Pagel’s lamda and Blomberg’s K test. (a) Distribution of Pagel’s obtained from 100 subsampling data with true bacterial origin vs data with randomised bacterial origins. (b) Distribution of Blomberg’s K obtained from 100 subsampled data with true and randomised bacterial origin. Elevated Pagel’s lamda and Blomberg’s K in true data compared to randomised data suggest that there are genetic signals associated with bacterial clinical and environmental origins. (c) Correlation between bacterial source (clinical or environmental) and tree with partial information. Log-likelihood (y-axis) was used to estimate the fitness of source following tree transformation by Pagel’s λ (x-axis). A multiplication to internal branches with λ = 0 created a basal tree, while a multiplication with λ = 1 kept the tree unchanged. d) Correlation between origin and tree when the origin was randomized (n=100). Three different probability models for trait transition were employed for both c) and d) including an equal rate model (ER), a symmetric model (SYM), and all rates different matrix (ARD).

**Supplementary Fig. 4.**
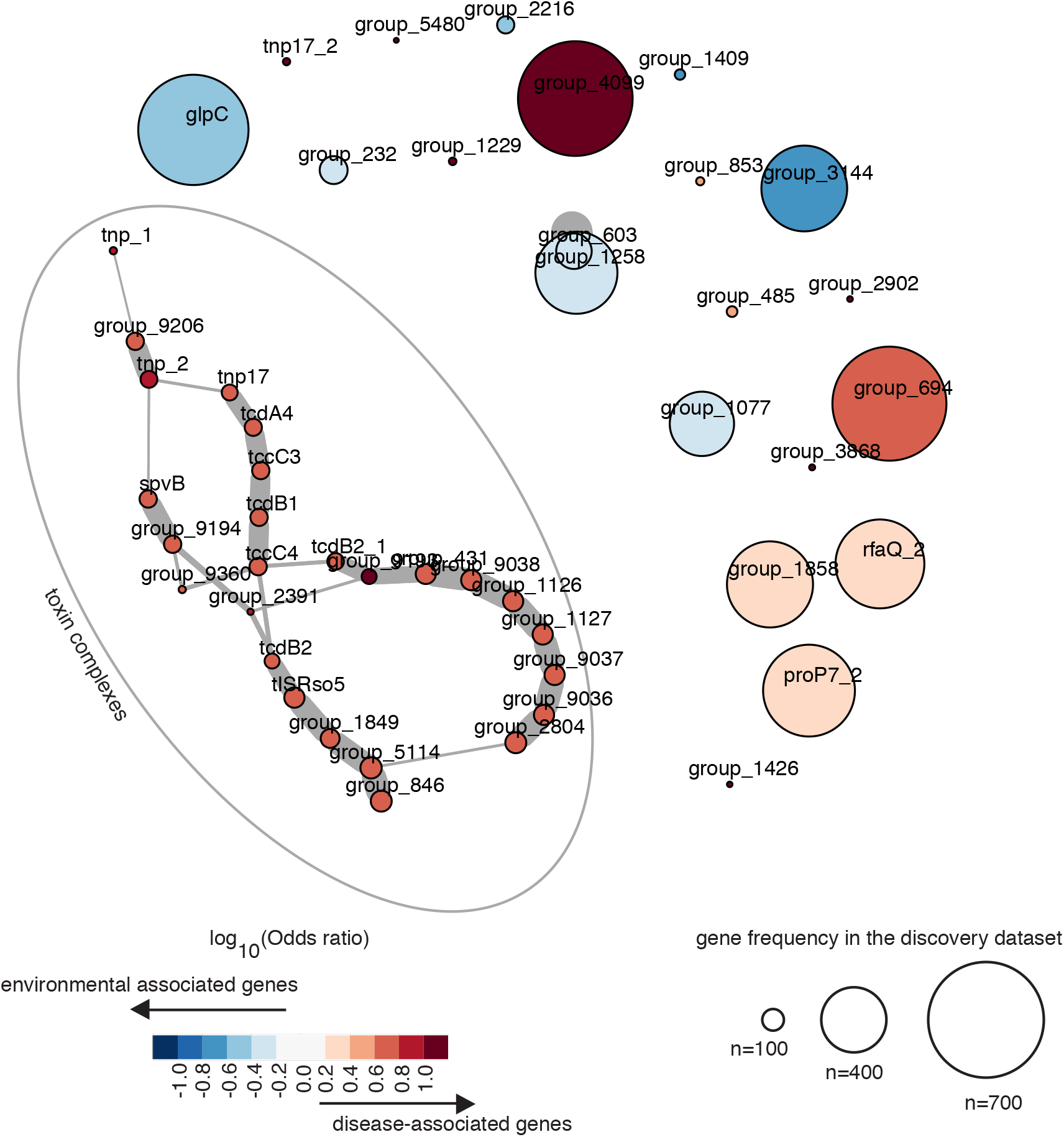
Distance network reveals genetic loci enriched in disease- and environment-associated isolates. A network was constructed on distance between disease and environmental-associated genes that fell within the size of operon described by the transcriptional unit, as reported in Ooi *et al.* 2013. Each node represents each gene, with the edge thickness proportional to the frequency of each gene pair observed in the population. The largest disease-associated locus identified in this dataset was the toxin complex. The node colour indicates the effect size and directionality of association on the scale of log10(0dds ratio), with red and blue presenting association with disease and the environment, respectively.

**Supplementary Fig. 5.**
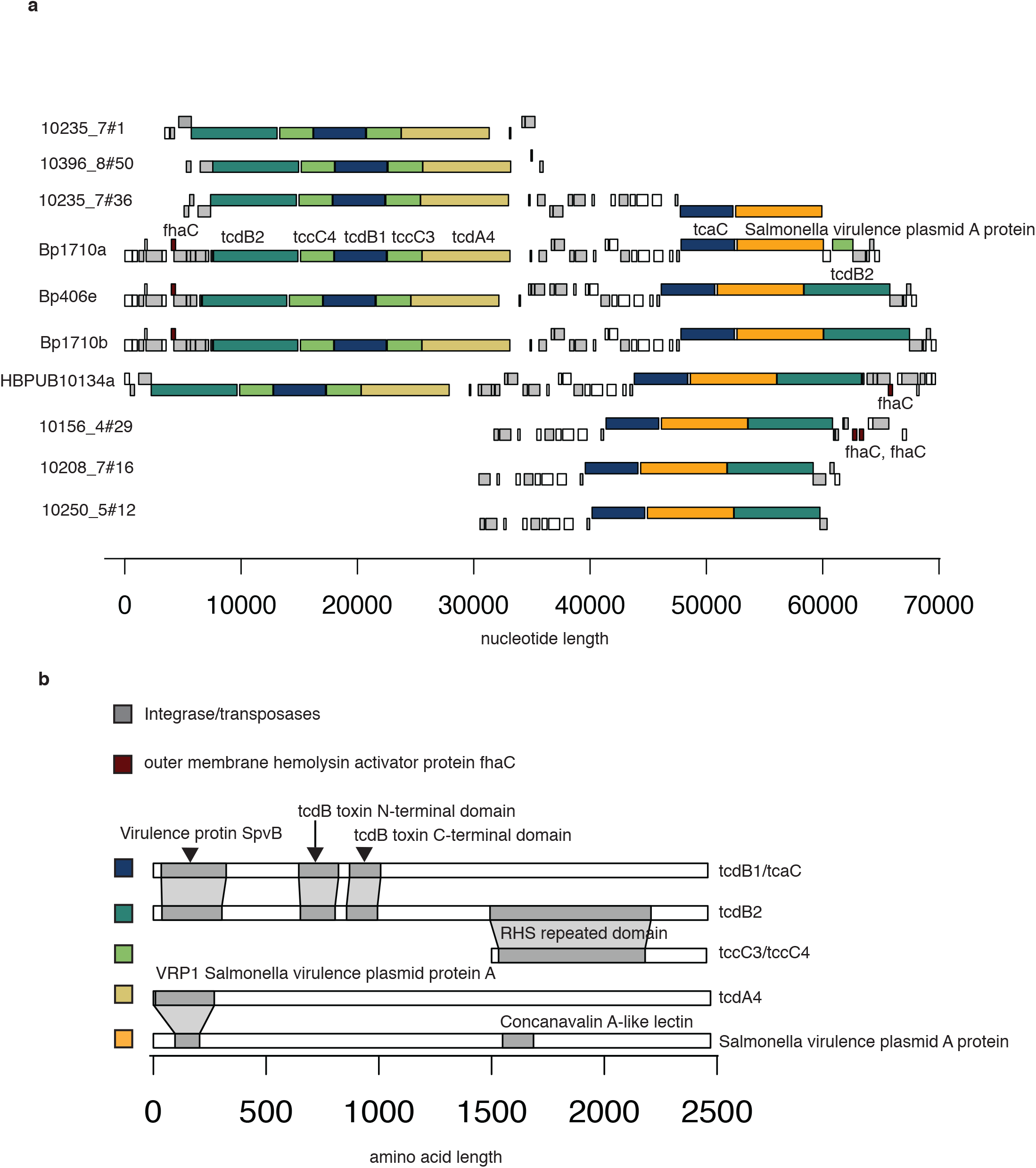
The genetic architecture of toxin-complexes, dominant hits comprising toxin genes that have been characterised previously in other bacterial species and transposable elements. (a) Gene order obtained from reference and newly assembled genomes from the discovery cohort. (b) Architecture of protein domains in each gene found in the toxin complex.

**Supplementary Fig. 6.**
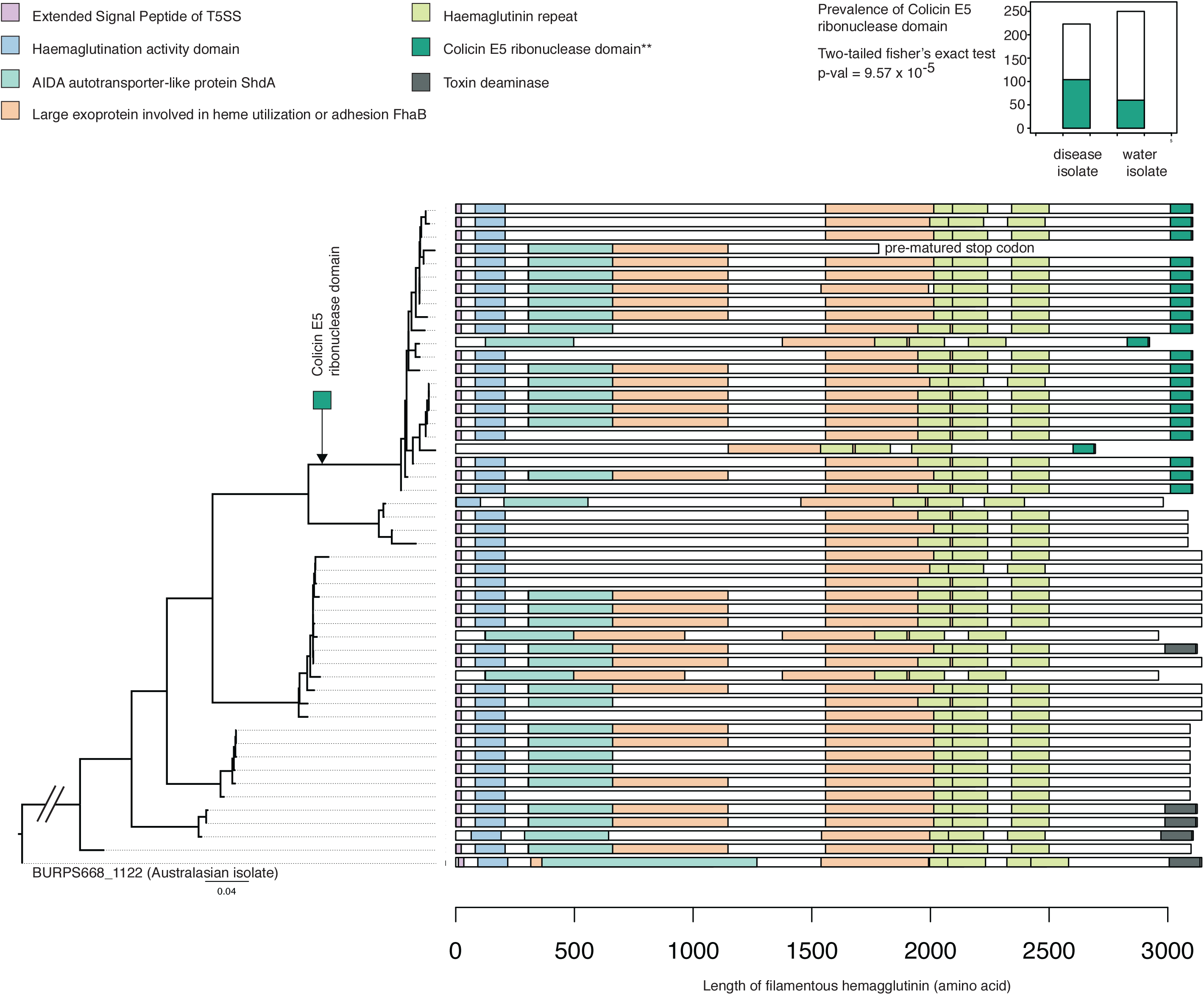
Architecture of protein domains of the filamentous hemagglutinin (BPSS2053), and the truncated form associated with the environmental isolates. A maximum likelihood phylogenetic tree (left) was constructed from 49 *fhaB* alleles found in the discovery dataset rooted on fhaB from an Australian outgroup (Bp668). Protein domains (right) were annotated onto each allele, highlighting the truncated form of filamentous hemagglutinin. The diagram also highlights loss of toxin deaminase, and gain of Colicin E5 ribonuclease domain at the C-terminus, respectively. The latter was present at an elevated frequency in clinical versus environmental isolates.

## References

1 Limmathurotsakul, D. et al. Predicted global distribution of Burkholderia pseudomallei and burden of melioidosis. Nat Microbiol 1, 15008, doi:10.1038/nmicrobiol.2015.8 (2016).

2 Wiersinga, W. J. et al. Melioidosis. Nat Rev Dis Primers 4, 17107, doi:10.1038/nrdp.2017.107 (2018).

3 Noinarin, P., Chareonsudjai, P., Wangsomnuk, P., Wongratanacheewin, S. & Chareonsudjai, S. Environmental Free-Living Amoebae Isolated from Soil in Khon Kaen, Thailand, Antagonize Burkholderia pseudomallei. PLoS One 11, e0167355, doi:10.1371/journal.pone.0167355 (2016).

4 Kaestli, M. et al. What drives the occurrence of the melioidosis bacterium Burkholderia pseudomallei in domestic gardens? PLoS Negl Trop Dis 9, e0003635, doi:10.1371/journal.pntd.0003635 (2015).

5 Limmathurotsakul, D. et al. Activities of daily living associated with acquisition of melioidosis in northeast Thailand: a matched case-control study. PLoS Negl Trop Dis 7, e2072, doi:10.1371/journal.pntd.0002072 (2013).

6 Currie, B. J., Ward, L. & Cheng, A. C. The epidemiology and clinical spectrum of melioidosis: 540 cases from the 20 year Darwin prospective study. PLoS Negl Trop Dis 4, e900, doi:10.1371/journal.pntd.0000900 (2010).

7 Holland, D. J., Wesley, A., Drinkovic, D. & Currie, B. J. Cystic Fibrosis and Burkholderia pseudomallei Infection: An Emerging Problem? Clin Infect Dis 35, e138–140, doi:10.1086/344447 (2002).

8 Ralph, A., McBride, J. & Currie, B. J. Transmission of Burkholderia pseudomallei via breast milk in northern Australia. Pediatr Infect Dis J 23, 1169–1171 (2004).

9 Wuthiekanun, V. et al. Development of antibodies to Burkholderia pseudomallei during childhood in melioidosis-endemic northeast Thailand. Am J Trop Med Hyg 74, 1074–1075 (2006).

10 Vasu, C., Vadivelu, J. & Puthucheary, S. D. The humoral immune response in melioidosis patients during therapy. Infection 31, 24–30, doi:10.1007/s15010-002-3020-2 (2003).

11 Teparrukkul, P. et al. Gastrointestinal tract involvement in melioidosis. Trans R Soc Trop Med Hyg 111, 185–187, doi:10.1093/trstmh/trx031 (2017).

12 Goodyear, A., Bielefeldt-Ohmann, H., Schweizer, H. & Dow, S. Persistent gastric colonization with Burkholderia pseudomallei and dissemination from the gastrointestinal tract following mucosal inoculation of mice. PLoS One 7, e37324, doi:10.1371/journal.pone.0037324 (2012).

13 Bragonzi, A. et al. Environmental Burkholderia cenocepacia Strain Enhances Fitness by Serial Passages during Long-Term Chronic Airways Infection in Mice. Int J Mol Sci 18, doi:10.3390/ijms18112417 (2017).

14 Massey, S. et al. Comparative Burkholderia pseudomallei natural history virulence studies using an aerosol murine model of infection. Sci Rep 4, 4305, doi:10.1038/srep04305 (2014).

15 Welkos, S. L. et al. Characterization of Burkholderia pseudomallei Strains Using a Murine Intraperitoneal Infection Model and In Vitro Macrophage Assays. PLoS One 10, e0124667, doi:10.1371/journal.pone.0124667 (2015).

16 Lewis, E. R. G., Kilgore, P. B., Mott, T. M., Pradenas, G. A. & Torres, A. G. Comparing in vitro and in vivo virulence phenotypes of Burkholderia pseudomallei type G strains. PLoS One 12, e0175983, doi:10.1371/journal.pone.0175983 (2017).

17 Sahl, J. W. et al. The Effects of Signal Erosion and Core Genome Reduction on the Identification of Diagnostic Markers. MBio 7, doi:10.1128/mBio.00846-16 (2016).

18 Spring-Pearson, S. M. et al. Pangenome Analysis of Burkholderia pseudomallei: Genome Evolution Preserves Gene Order despite High Recombination Rates. PLoS One 10, e0140274, doi:10.1371/journal.pone.0140274 (2015).

19 Chewapreecha, C. et al. Global and regional dissemination and evolution of Burkholderia pseudomallei. Nat Microbiol 2, 16263, doi:10.1038/nmicrobiol.2016.263 (2017).

20 Johnson, S. L. et al. Complete genome sequences for 59 burkholderia isolates, both pathogenic and near neighbor. Genome Announc 3, doi:10.1128/genomeA.00159-15 (2015).

21 Daligault, H. E. et al. Whole-genome assemblies of 56 burkholderia species. Genome Announc 2, doi:10.1128/genomeA.01106-14 (2014).

22 Viberg, L. T. et al. Whole-Genome Sequences of Five Burkholderia pseudomallei Isolates from Australian Cystic Fibrosis Patients. Genome Announc 3, doi:10.1128/genomeA.00254-15 (2015).

23 Price, E. P. et al. Unprecedented Melioidosis Cases in Northern Australia Caused by an Asian Burkholderia pseudomallei Strain Identified by Using Large-Scale Comparative Genomics. Appl Environ Microbiol 82, 954–963, doi:10.1128/AEM.03013-15 (2016).

24 Godoy, D. et al. Multilocus sequence typing and evolutionary relationships among the causative agents of melioidosis and glanders, Burkholderia pseudomallei and Burkholderia mallei. J Clin Microbiol 41, 2068–2079 (2003).

25 Price, E. P. et al. Improved multilocus sequence typing of Burkholderia pseudomallei and closely related species. J Med Microbiol 65, 992–997, doi:10.1099/jmm.0.000312 (2016).

26 Page, A. J. et al. Roary: rapid large-scale prokaryote pan genome analysis. Bioinformatics 31, 3691–3693, doi:10.1093/bioinformatics/btv421 (2015).

27 Lees, J. A. et al. Sequence element enrichment analysis to determine the genetic basis of bacterial phenotypes. Nat Commun 7, 12797, doi:10.1038/ncomms12797 (2016).

28 Limmathurotsakul, D. et al. Melioidosis caused by Burkholderia pseudomallei in drinking water, Thailand, 2012. Emerg Infect Dis 20, 265–268, doi:10.3201/eid2002.121891 (2014).

29 McRobb, E. et al. Distribution of Burkholderia pseudomallei in northern Australia, a land of diversity. Appl Environ Microbiol 80, 3463–3468, doi:10.1128/AEM.00128-14 (2014).

30 Viberg, L. T. et al. Within-Host Evolution of Burkholderia pseudomallei during Chronic Infection of Seven Australasian Cystic Fibrosis Patients. MBio 8, doi:10.1128/mBio.00356-17 (2017).

31 Nandi, T. et al. Burkholderia pseudomallei sequencing identifies genomic clades with distinct recombination, accessory, and epigenetic profiles. Genome Res 25, 608 (2015).

32 Currie, B. J. et al. Use of Whole-Genome Sequencing to Link Burkholderia pseudomallei from Air Sampling to Mediastinal Melioidosis, Australia. Emerg Infect Dis 21, 2052–2054, doi:10.3201/eid2111.141802 (2015).

33 Merhej, V., Georgiades, K. & Raoult, D. Postgenomic analysis of bacterial pathogens repertoire reveals genome reduction rather than virulence factors. Brief Funct Genomics 12, 291–304, doi:10.1093/bfgp/elt015 (2013).

34 Weinert, L. A. et al. Genomic signatures of human and animal disease in the zoonotic pathogen Streptococcus suis. Nat Commun 6, 6740, doi:10.1038/ncomms7740 (2015).

35 Pagel, M. Inferring the historical patterns of biological evolution. Nature 401, 877–884, doi:10.1038/44766 (1999).

36 Blomberg, S. P., Garland, T., Jr. & Ives, A. R. Testing for phylogenetic signal in comparative data: behavioral traits are more labile. Evolution 57, 717–745 (2003).

37 Geoghegan, J. L. & Holmes, E. C. The phylogenomics of evolving virus virulence. Nat Rev Genet, doi:10.1038/s41576-018-0055-5 (2018).

38 Brynildsrud, O., Bohlin, J., Scheffer, L. & Eldholm, V. Rapid scoring of genes in microbial pan-genome-wide association studies with Scoary. Genome Biol 17, 238, doi:10.1186/s13059-016-1108-8 (2016).

39 Ooi, W. F. et al. The condition-dependent transcriptional landscape of Burkholderia pseudomallei. PLoS Genet 9, e1003795, doi:10.1371/journal.pgen.1003795 (2013).

40 Price, E. P. et al. Unprecedented Melioidosis Cases in Northern Australia Caused by an Asian Burkholderia pseudomallei Strain Identified by Using Large-Scale Comparative Genomics. Appl Environ Microbiol 82, 954–963, doi:10.1128/AEM.03013-15 (2015).

41 Pearson, T. et al. Phylogeographic reconstruction of a bacterial species with high levels of lateral gene transfer. BMC Biol 7, 78, doi:10.1186/1741-7007-7-78 (2009).

42 Yang, G. & Waterfield, N. R. The role of TcdB and TccC subunits in secretion of the Photorhabdus Tcd toxin complex. PLoS Pathog 9, e1003644, doi:10.1371/journal.ppat.1003644 (2013).

43 Gatsogiannis, C. et al. A syringe-like injection mechanism in Photorhabdus luminescens toxins. Nature 495, 520–523, doi:10.1038/nature11987 (2013).

44 Chen, W. J. et al. Characterization of an insecticidal toxin and pathogenicity of Pseudomonas taiwanensis against insects. PLoS Pathog 10, e1004288, doi:10.1371/journal.ppat.1004288 (2014).

45 Sarovich, D. S. et al. Variable virulence factors in Burkholderia pseudomallei (melioidosis) associated with human disease. PLoS One 9, e91682, doi:10.1371/journal.pone.0091682 (2014).

46 Nandi, T. et al. A genomic survey of positive selection in Burkholderia pseudomallei provides insights into the evolution of accidental virulence. PLoS Pathog 6, e1000845, doi:10.1371/journal.ppat.1000845 (2010).

47 Brown, S. P., Cornforth, D. M. & Mideo, N. Evolution of virulence in opportunistic pathogens: generalism, plasticity, and control. Trends Microbiol 20, 336–342, doi:10.1016/j.tim.2012.04.005 (2012).

48 Limmathurotsakul, D. et al. Microevolution of Burkholderia pseudomallei during an acute infection. J Clin Microbiol 52, 3418–3421, doi:10.1128/JCM.01219-14 (2014).

49 Wood, D. E. & Salzberg, S. L. Kraken: ultrafast metagenomic sequence classification using exact alignments. Genome Biol 15, R46, doi:10.1186/gb-2014-15-3-r46 (2014).

50 Page, A. J. et al. Robust high-throughput prokaryote de novo assembly and improvement pipeline for Illumina data. Microb Genom 2, e000083, doi:10.1099/mgen.0.000083 (2016).

51 Seemann, T. Prokka: rapid prokaryotic genome annotation. Bioinformatics 30, 2068–2069, doi:10.1093/bioinformatics/btu153 (2014).

52 Holden, M. T. et al. Genomic plasticity of the causative agent of melioidosis, Burkholderia pseudomallei. Proc Natl Acad Sci U S A 101, 14240–14245, doi:10.1073/pnas.0403302101 (2004).

53 Ondov, B. D. et al. Mash: fast genome and metagenome distance estimation using MinHash. Genome Biol 17, 132, doi:10.1186/s13059-016-0997-x (2016).

54 Katoh, K. & Standley, D. M. MAFFT multiple sequence alignment software version 7: improvements in performance and usability. Mol Biol Evol 30, 772–780, doi:10.1093/molbev/mst010 (2013).

55 Galtier, N., Gouy, M. & Gautier, C. SEAVIEW and PHYLO_WIN: two graphic tools for sequence alignment and molecular phylogeny. Comput Appl Biosci 12, 543–548 (1996).

56 Page, A. J. et al. SNP-sites: rapid efficient extraction of SNPs from multi-FASTA alignments. Microbial Genomics 2, doi:doi:10.1099/mgen.0.000056 (2016).

57 Stamatakis, A. RAxML version 8: a tool for phylogenetic analysis and postanalysis of large phylogenies. Bioinformatics 30, 1312–1313, doi:10.1093/bioinformatics/btu033 (2014).

58 Molina-Venegas, R. & Rodriguez, M. A. Revisiting phylogenetic signal; strong or negligible impacts of polytomies and branch length information? BMC Evol Biol 17, 53, doi:10.1186/s12862-017-0898-y (2017).

59 Harmon, L. J., Weir, J. T., Brock, C. D., Glor, R. E. & Challenger, W. GEIGER: investigating evolutionary radiations. Bioinformatics 24, 129–131, doi:10.1093/bioinformatics/btm538 (2008).

60 Kembel, S. W. et al. Picante: R tools for integrating phylogenies and ecology. Bioinformatics 26, 1463–1464, doi:10.1093/bioinformatics/btq166 (2010).

61 Kent, W. J. BLAT--the BLAST-like alignment tool. Genome Res 12, 656–664, doi:10.1101/gr.229202. Article published online before March 2002 (2002).

62 Hunt, M. et al. ARIBA: rapid antimicrobial resistance genotyping directly from sequencing reads. Microb Genom 3, e000131, doi:10.1099/mgen.0.000131 (2017).

63 Finn, R. D. et al. InterPro in 2017-beyond protein family and domain annotations. Nucleic Acids Res 45, D190–D199, doi:10.1093/nar/gkw1107 (2017).

64 Winsor, G. L. et al. The Burkholderia Genome Database: facilitating flexible queries and comparative analyses. Bioinformatics 24, 2803–2804, doi:10.1093/bioinformatics/btn524 (2008).

65 Kanehisa, M., Furumichi, M., Tanabe, M., Sato, Y. & Morishima, K. KEGG: new perspectives on genomes, pathways, diseases and drugs. Nucleic Acids Res 45, D353–D361, doi:10.1093/nar/gkw1092 (2017).

66 Caspi, R. et al. The MetaCyc database of metabolic pathways and enzymes. Nucleic Acids Res 46, D633–D639, doi:10.1093/nar/gkx935 (2018).

67 Marchler-Bauer, A. et al. CDD/SPARCLE: functional classification of proteins via subfamily domain architectures. Nucleic Acids Res 45, D200–D203, doi:10.1093/nar/gkw1129 (2017).

68 Zhang, C., Wang, J., Long, M. & Fan, C. gKaKs: the pipeline for genome-level Ka/Ks calculation. Bioinformatics 29, 645–646, doi:10.1093/bioinformatics/btt009 (2013).

69 Revell, L. J. phytools: An R package for phylogenetic comparative biology (and other things). Methods Ecol. Evol. 3, 217–223 (2012).

70 Hadfield, J. et al. Phandango: an interactive viewer for bacterial population genomics. Bioinformatics, doi:10.1093/bioinformatics/btx610 (2017).

